# Parabricks: GPU Accelerated Universal Pan-Instrument Genomics Analysis Software Suite

**DOI:** 10.1101/2025.07.23.666378

**Authors:** Tong Zhu, Pankaj Vats, Seth Onken, Al Dunstan, Babak Zamirai, Daniel F. Puleri, Abhishek Nair, Marco Oliva, Anil Gaihre, Priyanka Sadhnani, Shaomeng Li, Kamesh Arumugam, Alejandro Chacon, Milos Maric, Jonathan Cohen, Ankit Sethia, Mehrzad Samadi

## Abstract

Next-generation sequencing (NGS) has transformed genomics, enabling breakthroughs in biotechnology, healthcare, and pharmaceuticals. However, exponential data growth-outpacing Moore’s Law-presents critical computational challenges for secondary analysis. We introduce Parabricks, a freely accessible, GPU-accelerated software suite supporting diverse workflows, including wholegenome, exome, transcriptome, and methylation analysis. Compatible across major short read and long read sequencing platforms, it reduces processing time from hours to minutes while maintaining concordance with established standards, this acceleration translates to significant cost savings. Deployable on-premises or through cloud platforms, Parabricks ensures scalable, genomic processing. Integrated AI-driven workflows and deep learning models further enhance precision, offering a universal solution for high-throughput genomic research and clinical diagnostics.

## 1 Main

The landscape of genome sequencing has advanced rapidly over the past decade, achieving remarkable milestones in both speed and cost-effectiveness. Current technological capabilities enable complete human genome sequencing within 24 hours at approximately $100 per genome [1–3]. This dramatic reduction in sequencing costs and time has fundamentally democratized access to Whole Genome Sequencing (WGS), along with Whole Exome Sequencing (WES), and Transcriptome Sequencing, fueling large-scale genomic initiatives and an exponential growth of Next-Generation Sequencing (NGS) data [2, 4]. Despite these technological advances in sequencing, the processing of increasingly voluminous genomic datasets presents great computational challenges that demand scalable, reproducible bioinformatics pipelines capable of maintaining pace with rapidly evolving sequencing technologies. Recent NGS workflows, which encompass critical processes such as sequence alignment, variant calling, genome assembly, and comprehensive downstream analyses, impose substantial computational resource requirements. This computational bottleneck has created an urgent need for innovative computing frameworks that can effectively accelerate and optimize genomic data processing across both research and clinical applications.

To address the increasing computational demands of modern genomics, Graphics Processing Units (GPUs) have emerged as a transformative solution, leveraging their parallel processing capabilities to dramatically accelerate genomic analyses [5–9]. In response to these challenges, we introduce NVIDIA Parabricks, a comprehensive, freely available GPU-accelerated software suite aimed to facilitate efficient and comprehensive genome analysis. As illustrated in Figure 1, NVIDIA Parabricks suite provides a comprehensive collection of GPU-accelerated genomics tools designed to enhance secondary genomic analysis workflows. Parabricks encompasses essential bioinformatics algorithms, from sequence alignment through variant calling and quality control, all optimized for high-performance computing environments. The complete workflow implementation is demonstrated through standardized pipeline frameworks, specifically WDL (Workflow Description Language) and Nextflow workflows, as detailed in Figure 6. https://docs.nvidia.com/clara/parabricks/4.5.0/documentation/tooldocs/wdlnextflow.html [5, 6, 9, 10]. Parabricks is compatible with all major next-generation sequencing (NGS) platforms, including both short-read and long-read technologies.

**Fig. 1.**
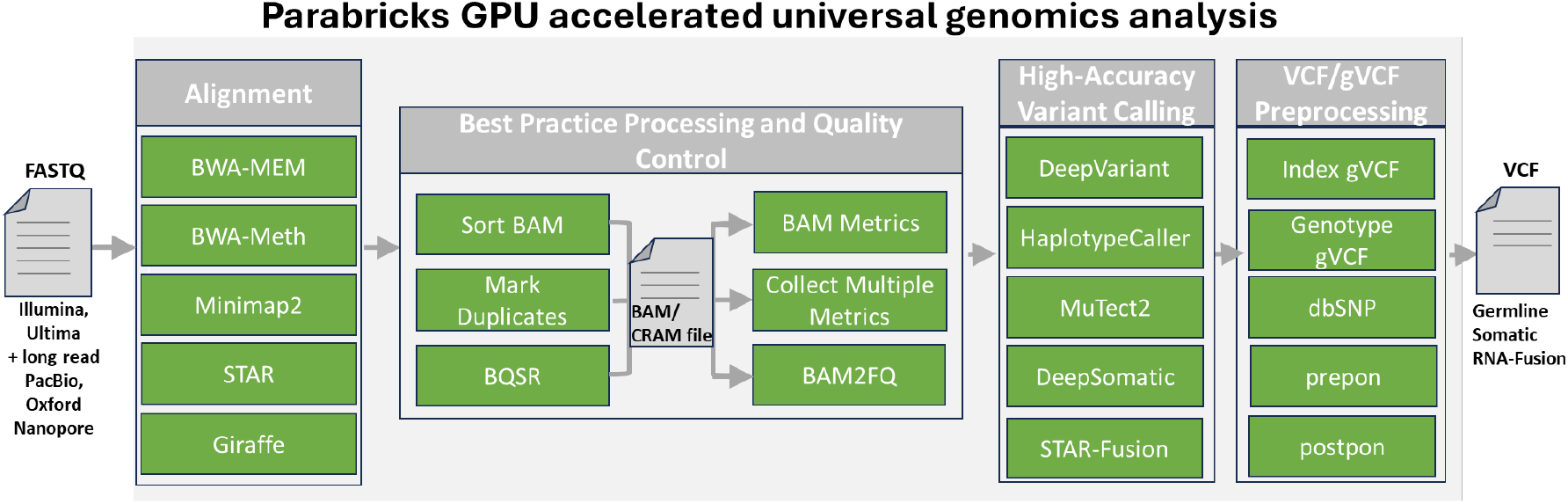
Overview of Parabricks the GPU accelerated universal genomics analysis software suite (can handle Illumina, Ultima, Singular, MGI, Element Biosciences, Pac-Bio and ONT dataset).Achieving up to 50% lower compute cost and 100x faster for WGS analysis compared to CPU only baseline. Provide Higher accuracy with option of retraining the DeepVariant on custom datasets.

Parabricks is designed to streamline and accelerate a comprehensive range of genomic analysis modules by integrating industry-standard aligners such as BWA-MEM [11], Minimap2 [12], and pangenome-aware Giraffe [13], as well as providing accelerated BWA-Meth [14] for bisulfite sequencing. Furthermore, the suite incorporates GPU-accelerated algorithms for essential post-processing steps, including BAM sorting, base quality score recalibration (BQSR), and duplicate read marking [15– 19]. In addition to these core functionalities, Parabricks offers comprehensive support for state-of-the-art variant callers, including HaplotypeCaller [20, 21], MuTect2 [22], DeepVariant [23], and DeepSomatic [24], thereby ensuring high-accuracy detection of both germline and somatic variants. Its GPU-accelerated workflows extend to RNA-seq analysis, leveraging tools such as STAR [25], STARsolo, and STAR-Fusion [26] for rapid and scalable transcriptomic data processing. Moreover, utility tools like bam2fq provide enhanced flexibility by enabling efficient conversion of BAM or CRAM files back to FASTQ format, thus facilitating streamlined re-alignment and downstream analysis.

Performance benchmarking demonstrates the substantial computational advantages delivered by Parabricks across multiple sequencing technologies. For short-read germline workflows, as illustrated in Figure 2, the platform achieves remarkable acceleration when deployed on GPU architectures. Specifically, utilizing 4×T4, 4×A100, or 4×H100 GPUs, Parabricks delivers 11× to 38× speedup relative to conventional CPU-based pipelines, reducing end-to-end processing time for a 35× whole genome sequencing (WGS) dataset from 15 hours to 24 minutes. Parabricks maintains exceptional performance across long-read sequencing technologies. As demonstrated in Figure 3, Parabricks achieves 2.3× to 8× acceleration on PacBio datasets, completing 35× WGS analysis within 33 minutes. Most notably, Figure 4 highlights the platform’s superior performance on Oxford Nanopore R10 chemistry data, where Parabricks achieves up to 50× acceleration, reducing DeepVariant analysis time to merely 13 minutes for a 30× WGS dataset.

**Fig. 2.**
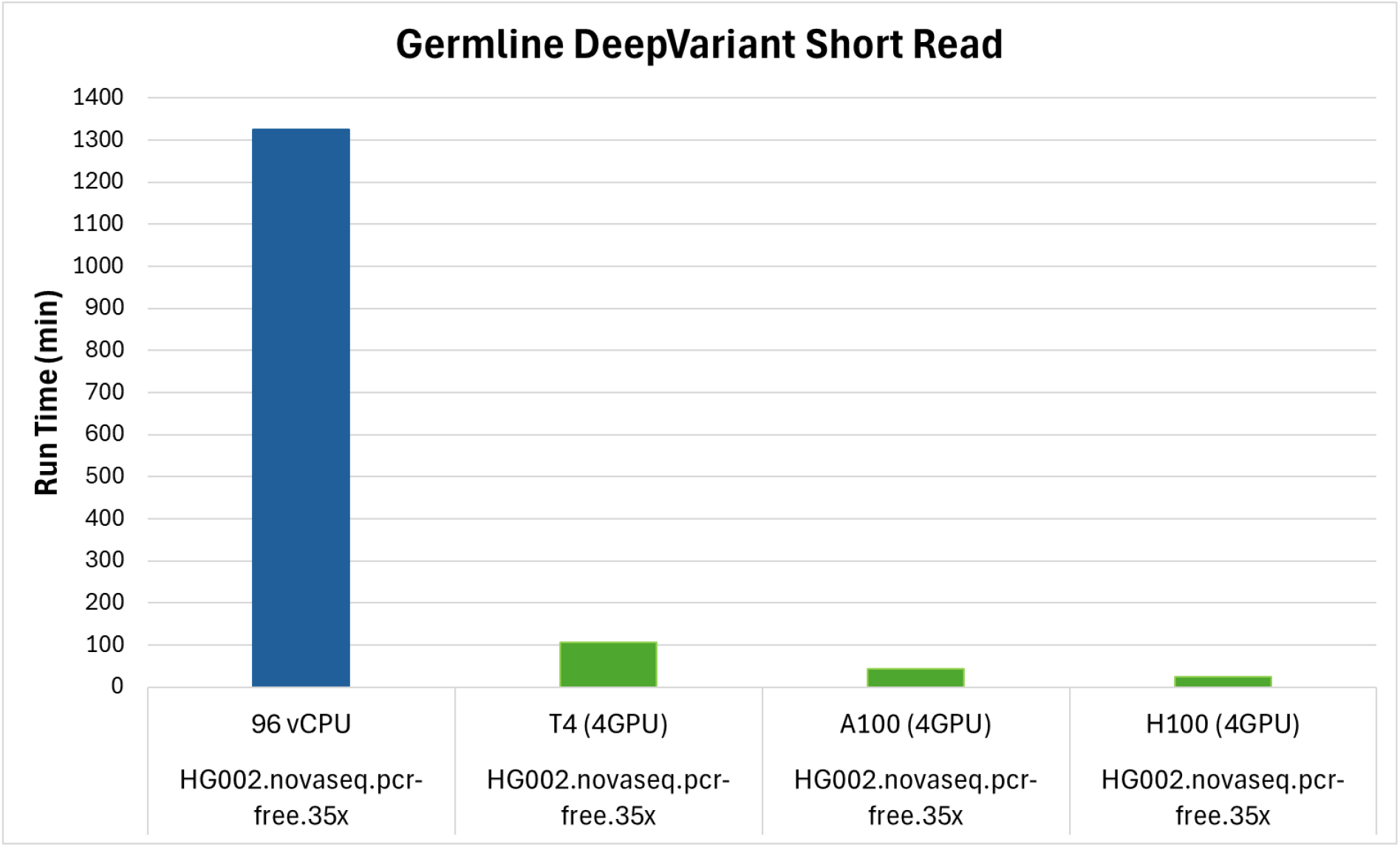
Acceleration of short-read germline variant calling workflows in Parabricks. Speedup comparison across GPU configurations (4×T4, 4×A100, 4×H100) versus 96-thread CPU baseline. Runtime breakdown for 35x short read WGS analysis showing alignment, sorting, duplicate marking, and variant calling stages shows 11x-38X acceleration on 4xT4, 4xA100, 4xH100 GPUs (right), resulting in a significant acceleration for GPu vs. CPU implementations.

**Fig. 3.**
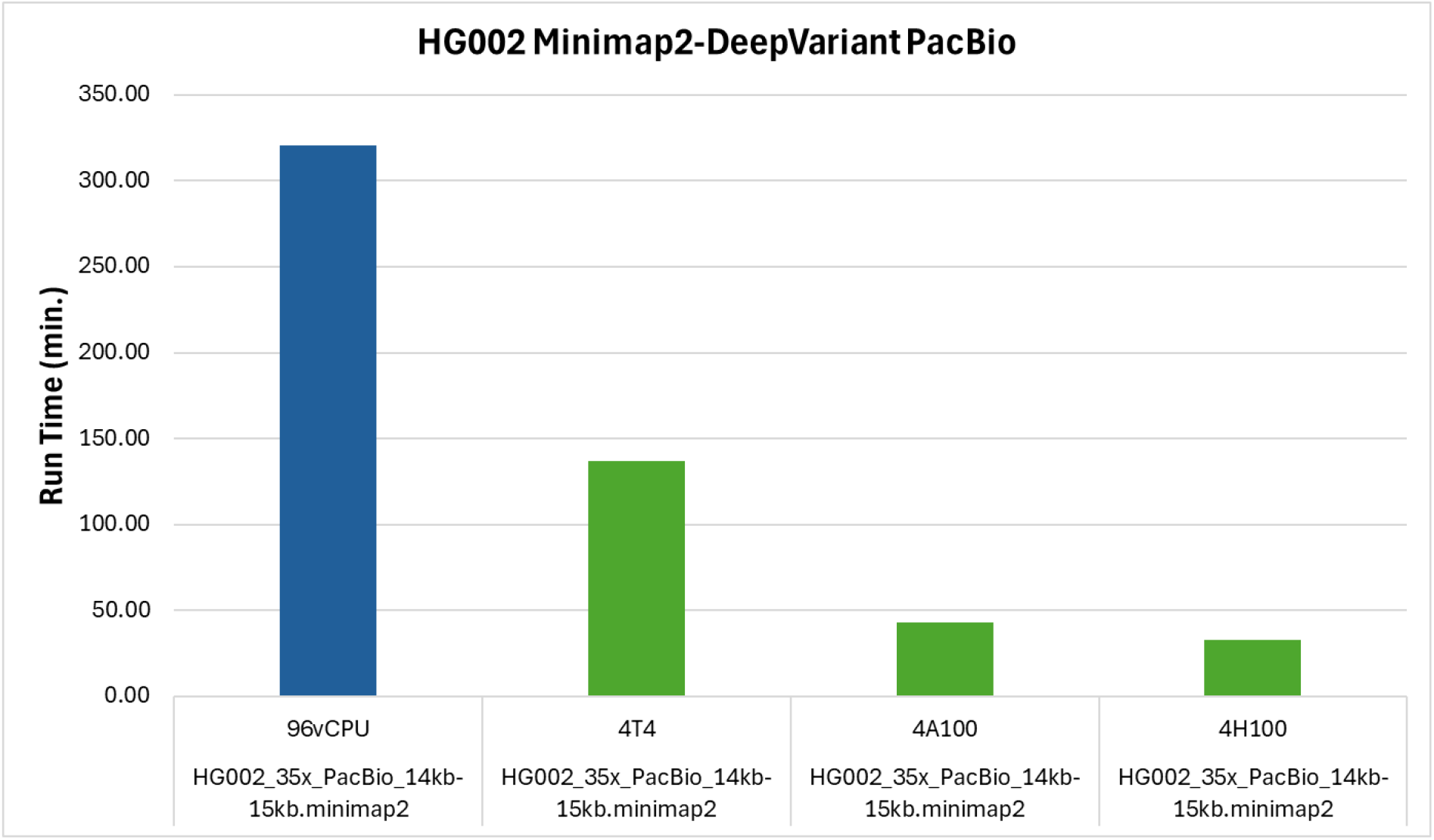
Acceleration of the Parabricks germline workflow on Pacbio dataset shows around 2.3x to 8x acceleration on 4xT4, 4xA100, 4xH100 GPUs (right), resulting in as little as 33 minutes end-to-end runtime for a 35x PacBio WGS sequencing dataset

**Fig. 4.**
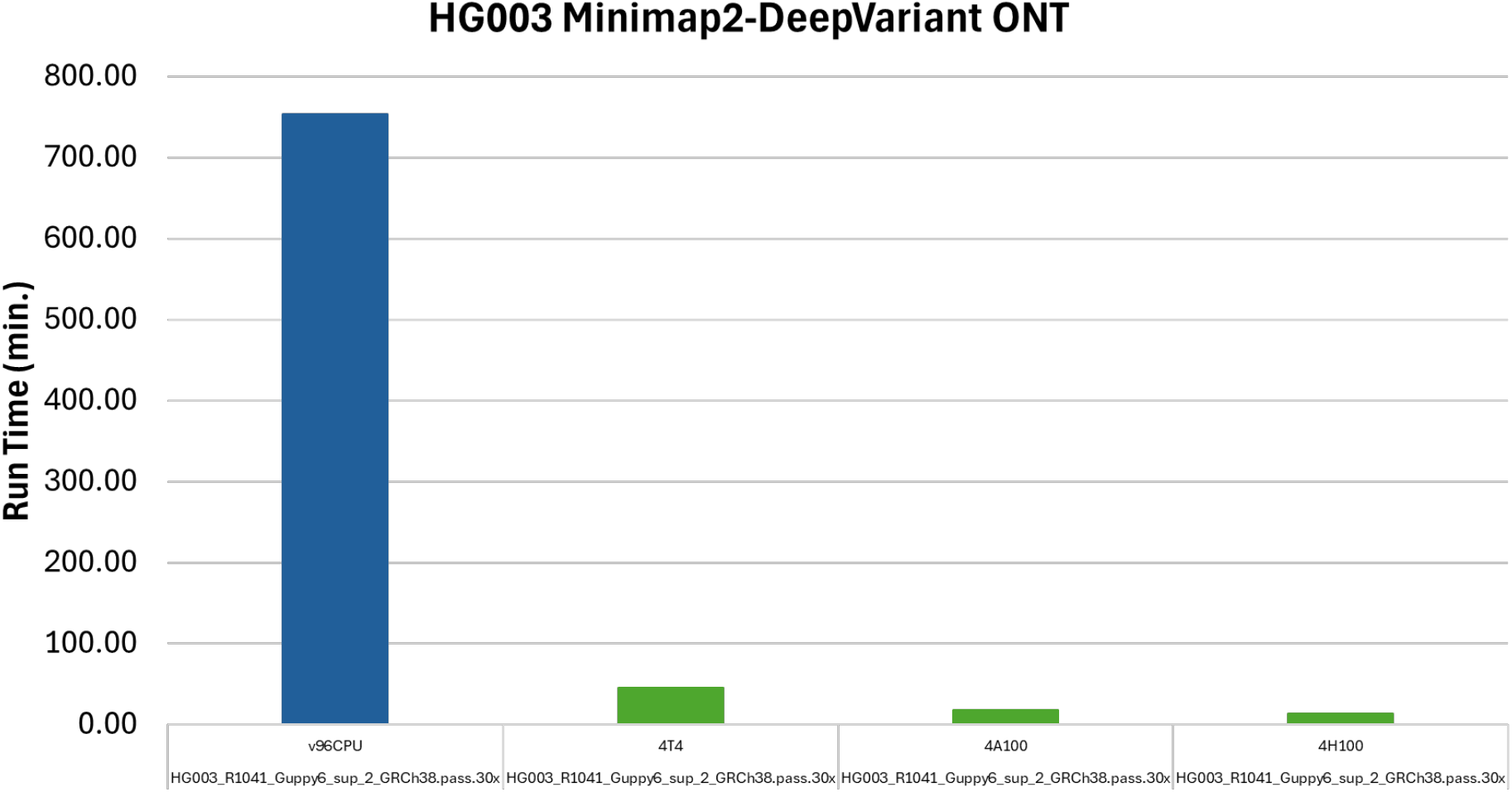
Acceleration of the Parabricks germline workflow on ONT R10 chemistry dataset shows up to 50x acceleration on 4xT4, 4xA100, 4xH100 GPUs (right), resulting in as little as 13 minutes for DeepVariant on a 35x ONT WGS sequencing dataset

Importantly, Parabricks achieves substantial speed improvements in genomic analysis without sacrificing accuracy. Across all supported platforms and workflows, it consistently maintains an F1 score exceeding 99.9% compared to baseline CPU-based tools, as illustrated in Figure 5, ensuring that rapid analysis is matched by clinical-grade variant calling performance. The combination of pan-platform acceleration and uncompromised accuracy not only accelerates the pace of genomic discovery but also makes high-throughput genomics more accessible and cost-effective for a broad range of scientific and clinical applications.

**Fig. 5.**
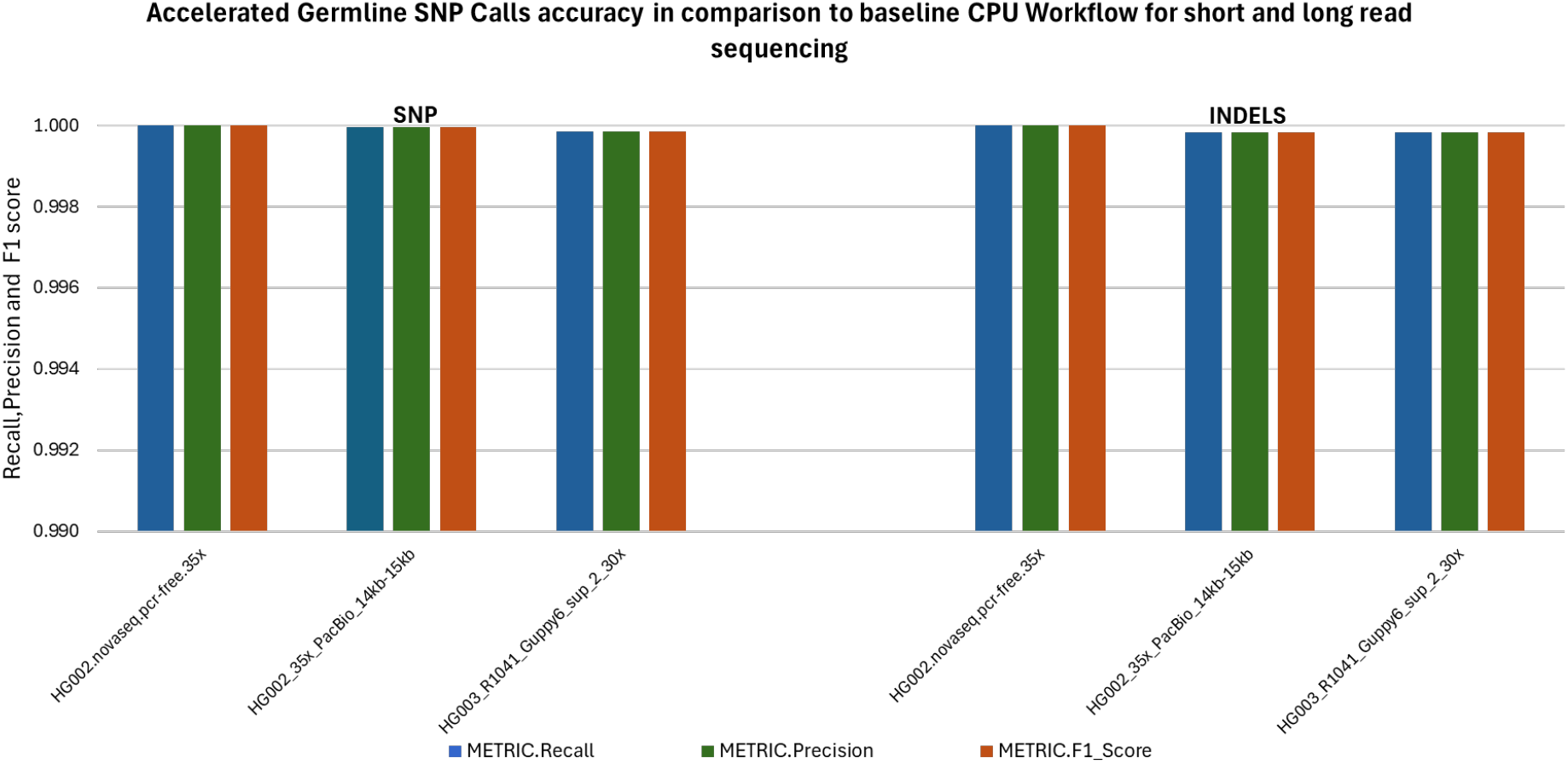
Parabricks accuracy benchmark for DeepVariant end to end germline workflow for short and long reads in comparison to base line CPU algorithms, Parabricks is able to reproduce the results with ¿99.9% Recall, Precision and F1 Score for both Short and Long read technologies.

**Fig. 6.**
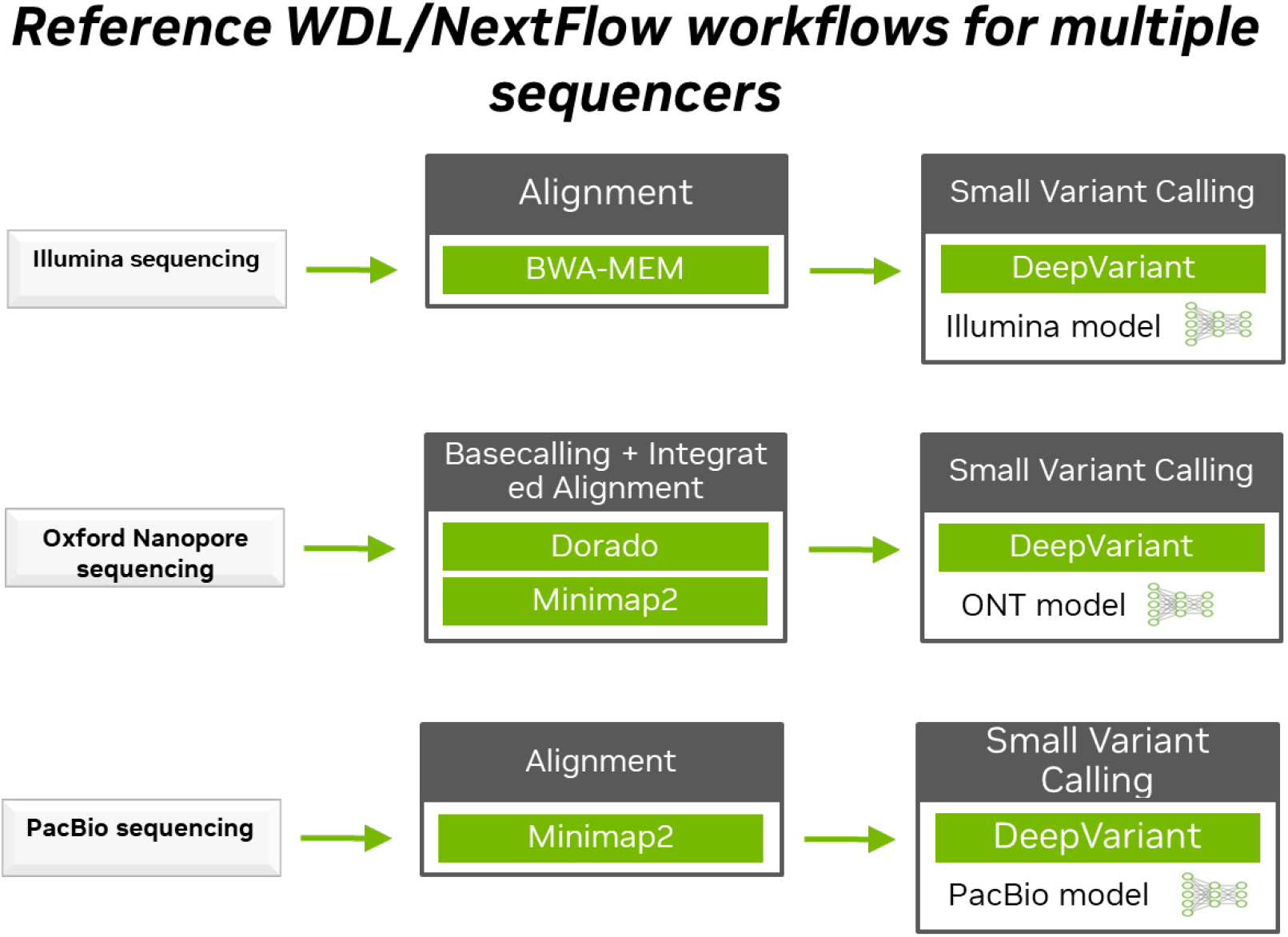
Parabricks containers are compatible with WDL and NextFlow for building customized workflows, here we are presenting the accelerated germline workflow for 3 different sequencing platform and can be deployed at scale.

The evolution of variant calling methodologies has been fundamentally transformed by deep learning innovations, which effectively address the inherent complexities of genomic data analysis. Traditional variant calling approaches often struggle with confounding factors including sequencing artifacts, alignment errors, and PCR-induced biases. Over the past decade, the genomics community has witnessed the emergence of diverse variant calling tools employing methodologies spanning from heuristic and statistical models such as Bayesian inference and Gaussian mixture models to sophisticated deep learning architectures incorporating convolutional and recurrent neural networks. Among these technological advances, DeepVariant stands out for its exceptional accuracy and broad generalizability across diverse sequencing technologies, genome builds, and mammalian species, as demonstrated by its leading performance in the Precision FDA Truth Challenge V2 [27], where deep learning-based methods dominated the field. DeepVariant’s success underscores the ability of deep learning approaches to generalize across different genome builds and mammalian species, adapting seamlessly to a broad range of sequencing technologies from short reads to long reads.

Building on this foundation, Parabricks further amplifies these advancements by integrating GPU-accelerated implementations of DeepVariant and DeepSomatic [23, 24], each optimized with parallel programming frameworks such as CUDA (Compute Unified Device Architecture) and TensorRT. These technical enhancements, combined with advanced retraining tools, enable researchers to fine tune models for specific laboratory protocols, sequencing platforms, and reagents. This integration not only ensures rapid variant analysis but also consistently delivers state-of-the-art accuracy, outperforming conventional CPU-based tools and underscoring the transformative potential of combining hardware acceleration with deep learning in precision genomics.

The practical significance of these technological advancements is evident in largescale, real-world deployments. For example, researchers at the Regeneron Genetics Center, in collaboration with UK Biobank and NVIDIA, leveraged Parabricks’ Deep-Variant retraining capabilities to sequence and analyze 500,000 exomes, achieving greater accuracy in genetic variant identification than traditional bioinformatics tools [28]. Similarly, the Wellcome Sanger Institute utilizes NVIDIA’s accelerated computing to enhance cancer genome analysis, significantly reducing costs and energy consumption [29].

By replacing traditional CPU-based workflows, Parabricks not only preserves output accuracy and configurability but also significantly enhances processing speed and reduces per-sample analysis costs. Importantly, Parabricks is designed to ensure results that are equivalent-and in most cases, identical-to those obtained using traditional CPU-based methods. When benchmarked against standard CPU pipelines for germline and somatic variant calling, Parabricks achieves an F1 score of 0.999 for both SNPs and indels, while simultaneously reducing execution time and computational costs. Any minor differences in output files can be attributed to factors such as pseudo-random number generation, rounding errors, library dependencies, or the use of additional arguments. This seamless combination of deep learning flexibility and GPU acceleration underscores Parabricks’ transformative potential in precision genomics, establishing it as a versatile, high-performance toolkit for a broad spectrum of genomic analyses [10].

The significance and effectiveness of Parabricks have been highlighted in numerous published case studies and collaborative projects [2, 4, 5, 9, 10, 30, 31], further validating its efficacy in processing data from extensive NGS platforms and its adaptability to a wide range of genomic analysis tasks. This versatility is especially valuable in fields where rapid and reliable genomic analysis is essential, such as public health, clinical management, and time-sensitive research areas like Neonatal Intensive Care Units (NICU) [2, 4] and neurodegenerative disease studies [9]. Parabricks genome analysis suite emerges as an efficient solution for developers of high-throughput sequencing centers, research labs and companies, capable of delivering seamless, real-time data analysis that keeps pace with data generation [32].

Parabricks leverages advanced NVIDIA technologies to address not only computational bottlenecks but also challenges related to data I/O and multiGPU utilization. It incorporates NVCOMP, a GPU-based compression library for fast, lossless data compression and decompression; NVIDIA TensorRT, an SDK for high-performance deep learning inference; GPUDirect Storage (GDS) for rapid, low-latency storage access; and Dynamic Programming X (DPX) instructions to accelerate dynamic programming algorithms commonly used in genomics and protein structure analysis. With the integration of these innovative technologies, Parabricks is freely available via NVIDIA GPU Cloud (NGC) (https://catalog.ngc.nvidia.com/orgs/nvidia/teams/clara/containers/claraparabricks), enabling easy deployment through Docker containers on any NVIDIA-certified system-on-premises or in the cloud. Comprehensive documentation and user guides are available (https://docs.nvidia.com/clara/parabricks/latest/index.html), and for those requiring enterprise-level support, a licensed version is offered through NVIDIA AI Enterprise.

In summary, Parabricks provides a free, pan-instrument, scalable, and user-friendly genomics software suite that harnesses GPU acceleration, advanced deep learning models, and refined genomics algorithms to deliver rapid, accurate, and cost-effective genomic analysis for the modern era.

## 2 Method

Acceleration of genomics analysis packages is achieved through efficient utilization of GPUs along with advanced NVIDIA technologies like CUDA DPX, NVCOMP, GDS, TensorRT inside Parabricks software suite [6–8]. On top of these techniques several modern computer software techniques including acceleration on GPUs, software pipelining, task level parallelism and overlapping of computation and data transfers have been applied extensively to reduce their wall-clock time. The fusion of these highperformance computing techniques with state-of-the-art genomics packages results in analysis that is not only faster but generates high quality results [10].

### 2.1 Germline variant

Germline variant calling was conducted using the well-characterized 35x Ashkenazim Trio (HG002 novaseq pcr free 35x) whole genome sequencing (WGS) dataset from the Genome in a Bottle (GIAB) consortium [33]. To establish baseline performance metrics, we implemented a standard CPU-based pipeline comprising BWA-MEM v0.7.15 [11] for read alignment and Deepvariant 1.6 [23] for germline variant calling. This standard workflow encompassed sequential steps of read alignment, sorting, duplicate marking, and subsequent variant calling using DeepVariant.

To rigorously assess computational efficiency and variant calling accuracy, we compared the performance of this conventional CPU workflow—executed on a highperformance server with 96 CPU threads—against the GPU-accelerated Parabricks germline pipeline. The Parabricks pipeline was systematically evaluated across multiple GPU configurations, including systems equipped with four NVIDIA T4, four A100, and four H100 GPUs. This comprehensive approach facilitated direct comparison of runtime, throughput, and variant calling concordance between CPU-based and GPU-accelerated methodologies.

To ensure comprehensive evaluation across diverse sequencing technologies, we extended our analysis to include long-read sequencing platforms. For long-read germline workflow analysis, we implemented a standard CPU-based pipeline comprising Minimap2 [12] for read alignment and Deepvariant 1.6 for variant calling. We processed both PacBio (HG002 35x PacBio 14kb-15kb) and Oxford Nanopore Technologies (ONT, HG003 R1041 Guppy6 30x) datasets. This approach enabled direct comparison of performance metrics across CPU based long-read germline variant calling vs GPU accelerated long read germline variant calling while maintaining consistent analytical parameters across all platforms.

### 2.2 DPX instruction

NVIDIA’s Hopper architecture introduced Dynamic Programming (DPX) instructions for the purpose of accelerating dynamic programming algorithms used in high performance computing. Of interest to Parabricks, is the Striped and banded SmithWaterman [34] algorithm, an algorithm introduced by Smith and Waterman for identifying homologous subregions between sequences of protein or DNA. The SmithWaterman algorithm is a computationally intensive process that constitutes a crucial part of BWA-MEM (accelerated in the Parabricks fq2bam tool). On Hopper GPUs, dedicated hardware support for DPX instructions allows multiple common dynamic programming operations to be done at the same time with a single instruction. These fused operations allow for higher throughput providing acceleration to DP algorithms. Parabricks leverages this to increase the Smith-Waterman throughput by a factor of 7.8x speedup on H100 over A100.

### 2.3 NVCOMP

NVIDIA nvCOMP is a library designed to accelerate data compression and decompression using GPUs. The library provides an interface to both compress and decompress data in the DEFLATE format [35], among others. Data is processed in parallel chunks; nvCOMP DEFLATE algorithm 1 achieves a compression throughput of 9.39 GB/s and decompression throughput of 18.19 GB/s on NVIDIA’s H100. Parabricks fq2bam uses nvCOMP in the GPU write mode to compress the final BAM (Binary Alignment Map) file on-device following alignment and sorting. The BAM file is a compressed binary file containing aligned reads in a gzip compatible BGZF format composed of DEFLATE compressed blocks which can be efficiently accessed.

### 2.4 GPUDirect Storage (GDS)

NVIDIA GPUDirect Storage (GDS) allows for the direct transfer of data between device memory and on-node storage, bypassing the traditional path through the host processor. GDS reduces the load on the CPU and allows data which has already been processed on GPU to be output to disk without an additional stop in host memory. Parabricks leverages this technology to accelerate I/O-bound parts of the application, such as the final steps of fq2bam. Using GDS allows the final preparation of alignments in a BAM format to be done entirely on GPU and transferred to disk, freeing up the CPU for other compute-intensive tasks and accelerating file output.

### 2.5 TensorRT

NVIDIA TensorRT, an SDK for high-performance deep learning inference, includes a deep learning inference optimizer and runtime that delivers low latency and high throughput for inference applications. TensorRT [36] is built on the NVIDIA CUDA (Compute Unified Device Architecture) parallel programming model and can perform TensorRT-based applications up to 36X faster than CPU-only platforms during inference, TensorRT performs different types of optimizations for increasing throughput of deep learning models, which includes Weight Activation Precision Calibration, Layer Tensor Fusion, Kernel Auto-Tuning, Dynamic Tensor Memory, Multi-Stream Execution, Time Fusion. Enable to optimize neural network models trained on all major frameworks, calibration for lower precision with high accuracy. Parabricks Deepvariant uses TensorRT to perform inference on pileup images, which requires a TensorRT engine file, and can be converted from the original Deepvariant TensorFlow checkpoint or saved model.

### 2.6 Computing environment and resources

Benchmarking for runtime was conducted using a variety of hardware configurations, including Amazon Web Services (AWS) instances as well as on-prem systems. For germline variant calling, we utilized the Ashkenazi Trio GIAB 35x whole genome dataset, We compared turnaround times and computational costs across several platforms: 4×T4 GPU nodes (g4dn.12xlarge AWS instances), on-premises systems using 4×A100 GPUs (DGX A100 320GB system containing 2×AMD EPYC 7742 and 8×A100-SXM4-40GB) and using 4×H100 GPUs (DGX H100 system containing 2×Intel Xeon Platinum 8480C and 8×H100 80GB HBM3), and traditional CPU-based processing using 96 threads on an m5.24xlarge AWS instance, following standard germline variant calling practice.

## Notes

https://catalog.ngc.nvidia.com/orgs/nvidia/teams/clara/containers/clara-parabricks

https://docs.nvidia.com/clara/parabricks/latest/index.html

https://docs.nvidia.com/clara/parabricks/4.5.0/documentation/tooldocs/wdlnextflow.html

### Competing Interest Statement

All the authors are salaried employees of NVIDIA, a leading technology company renowned for its pioneering work in graphics processing units (GPUs), artificial intelligence (AI), and high-performance computing.

### Summary of Updates

revised; author name correction and added ORCID id fixed link error on page 3. https://docs.nvidia.com/clara/parabricks/4.5.0/documentation/tooldocs/wdlnextflow.html

